# Deep learning of quality control for stereotaxic registration of human brain MRI

**DOI:** 10.1101/303487

**Authors:** Vladimir S. Fonov, Mahsa Dadar, The PREVENT-AD Research Group, D. Louis Collins

## Abstract

Linear registration to stereotaxic space is a common first step in many automated image-processing tools for analysis of human brain MRI scans. This step is crucial for the success of the following image-processing steps. Several well-established algorithms are commonly used in the field of neuroimaging for this task, but none of them has a 100% success rate. Manual assessment of the registration is commonly used as part of quality control.

We propose a completely automatic quality control method based on deep learning that replaces human rater and accurately performs quality control assessment for stereotaxic registration of T1w brain scans.

In a recently published study from our group comparing linear registration methods, we used a database of 9693 MRI scans from several publically available datasets and applied five linear registration tools. In this study, the resulting images that were assessed and labeled by a human rater are used to train a deep neural network to detect cases when registration failed.

Our method was able to achieve 88% accuracy and 11% false positive rate in detecting scans that should pass quality control, better than a manual QC rater.

## 1 Introduction

Many automatic image-processing techniques for processing human brain MRI scans include linear registration to stereotaxic space as one of the first steps in the pipeline [1, 2, 3]. Often, a human rater must manually verify the quality of this step by looking at a series of images that show overlap between the registered scan and some kind of reference template in order to identify datasets that failed registration. Such datasets may be subject to subsequent manual registration or may be discarded from analysis. For example, Ashburner et al. [4] explicitly states that quality control should be maintained as high as possible. Depending on the dataset and registration algorithm used, the success rate can vary from 100% down to 60% [5]. In particular, MRI scans of subjects with strong atrophy due to neurodegenerative diseases show lower rates of success when using registration tools that were originally developed and tested on MRI scans of young healthy subjects. Also, our experiments show that strong atrophy makes the task of manual quality control even more challenging when the reference template is representative of a healthy population.

As part of recently published work [20], to compare reliability of linear registration we performed an experiment where a large number of scans (9693 scans) from the Human Connectome Project (HCP) [14], the Alzheimer’s Disease Neuroimaging Initiative (ADNI) [12], the Pre-symptomatic evaluation of experimental or novel treatments for Alzheimer’s Disease (PreventAD) and the Parkinson’s Progression Marker Initiative (PPMI) [13] databases were linearly registered to the MNI-ICBM152 2009c space using five publically available linear registration methods. All registrations were then assessed and labeled by a human rater for quality. Since manual quality control of registrations is time consuming (∼30 hours for 9693 scans) and prone to inter-rater and intra-rater errors (intra-rater Dice index of 0.96), an automatic method that would be able to assess quality of registrations would be useful for the community. We attempted to replicate the behavior of a human rater in this task by training deep neural network (DNN) to determine quality of registrations by analyzing series of 2D control images.

Following the logic from [4], we determined that it is more important to ensure the quality of the images that were accepted by the rater, rather then overall accuracy of the classification, thus we decided to minimize False Positive Rate (FPR) metric (reflecting the proportion of incorrect registrations that were passed by the rater).

In order to leverage existing network design and speed training of the DNN, we adapted a pre-trained network that behaved well on image classification tasks [5]. In particular, we performed experiments using modified Resnet-18 (r18) [7, 11] and batch-normalized Network-In-Network (nin) [6, 10]. In our experiments, the resnet-18 was able to achieve performance similar to that of the human rater (in terms of accuracy) after 1500 iterations.

## 2 Methods and materials

### 2.1 Materials

We used T1w MRI scans from four different datasets (9693 scans in total):

- **ADNI**: The Alzheimer’s Disease Neuroimaging Initiative (ADNI) [12], is a multi-center and multi-scanner study with the aim of defining the progression of Alzheimer’s disease (AD). Subjects are normal controls, individuals with mild cognitive impairment or AD aged 55 years or older. Data was acquired using 1.5T and 3T scanners of different models of GE Medical Systems, Philips Medical systems, and SIEMENS at over 59 acquisition sites. We used 3489 scans from 1.5T scanners and 3056 from 3.0T.
- **PPMI**: The Parkinson Progression Marker Initiative (PPMI) [13] is an observational, multi-center and multi-scanner longitudinal study designed to identify PD biomarkers. Subjects are normal controls or de Novo Parkinson’s patients aged 30 years or older. Data was acquired using 1.5T and 3T scanners of different models of GE Medical Systems, Philips Medical systems, and SIEMENS at over 33 sites in 11 countries. We used 222 scans from 1.5T scanners and 778 from 3T.
- **HCP**: The Human Connectome Project (HCP) [14] is an effort to characterize brain connectivity and function and their variability in young healthy adults aged between 25 and 30 years. We used 897 scans.
- **PREVENT-AD**: The PREVENT-AD (Pre-symptomatic Evaluation of Novel or Experimental Treatments for Alzheimer’s Disease, http://www.prevent-alzheimer.ca) program [15] follows healthy individuals age 55 or older with a parental history of AD dementia. We used 1251 scans.

### 2.2 Automatic registration methods

We used the registration techniques briefly described below; a subset of the techniques used in [20]. For a full description of the methods see [20]. All available scans were registered to the stereotaxic space defined by MNI-ICBM152-2009c template (MNI template) [8]. All methods with the exception of Elastix used 9-parameters linear registration.

- **MRITOTAL**: a hierarchical multi-scale 3D registration technique for the purpose of aligning a given MRI volume to an average MRI template aligned with the Talairach stereotaxic coordinate system [16]. We have tested two configurations of this method: “standard” and “icbm”. The source code is available at https://github.com/BIC-MNI/mni_autoreg.
- **BestLinReg**: a 5-stage hierarchical technique similar to MRITOTAL that is part of the MINC tools and is based on a hierarchical non-linear registration strategy developed by Robbins et al. [17]. We tested two versions: one where cross-correlation coefficient is used as a cost function and another one using normalized mutual information. The source code is avaliable at https://github.com/BIC-MNI/EZminc/blob/ITK4/scripts/bestlinreg_s.
- **Revised BestLinReg**: This is the same as BestLinReg from above with different set of parameters, and normalized mutual information cost function. The source code is available at https://github.com/BIC-MNI/EZminc/blob/ITK4/scripts/bestlinreg_claude.pl.
- **Elastix**: an intensity-based registration tool [19]. Elastix has a parametric and modular framework, where the user can configure different components of the registration. We used Mattes mutual information, adaptive stochastic gradient descent optimizer with Similarity Transform with 7 parameters.

### 2.3 Manual quality control method

Our registration experiments produced 57848 linear registrations in total (some registration experiments crashed without producing usable results). Resulting transformations were stored as affine 4×4 matrices and applied to the corresponding scans to resample them on a 1mm^3^ voxel grid in stereotaxic space. Then, a series of slices were extracted, and the outline of the MNI template brain was overlaid on top to create an image that was given to the human rater to assess registration performance (See Figure 1).

**Figure 1.**
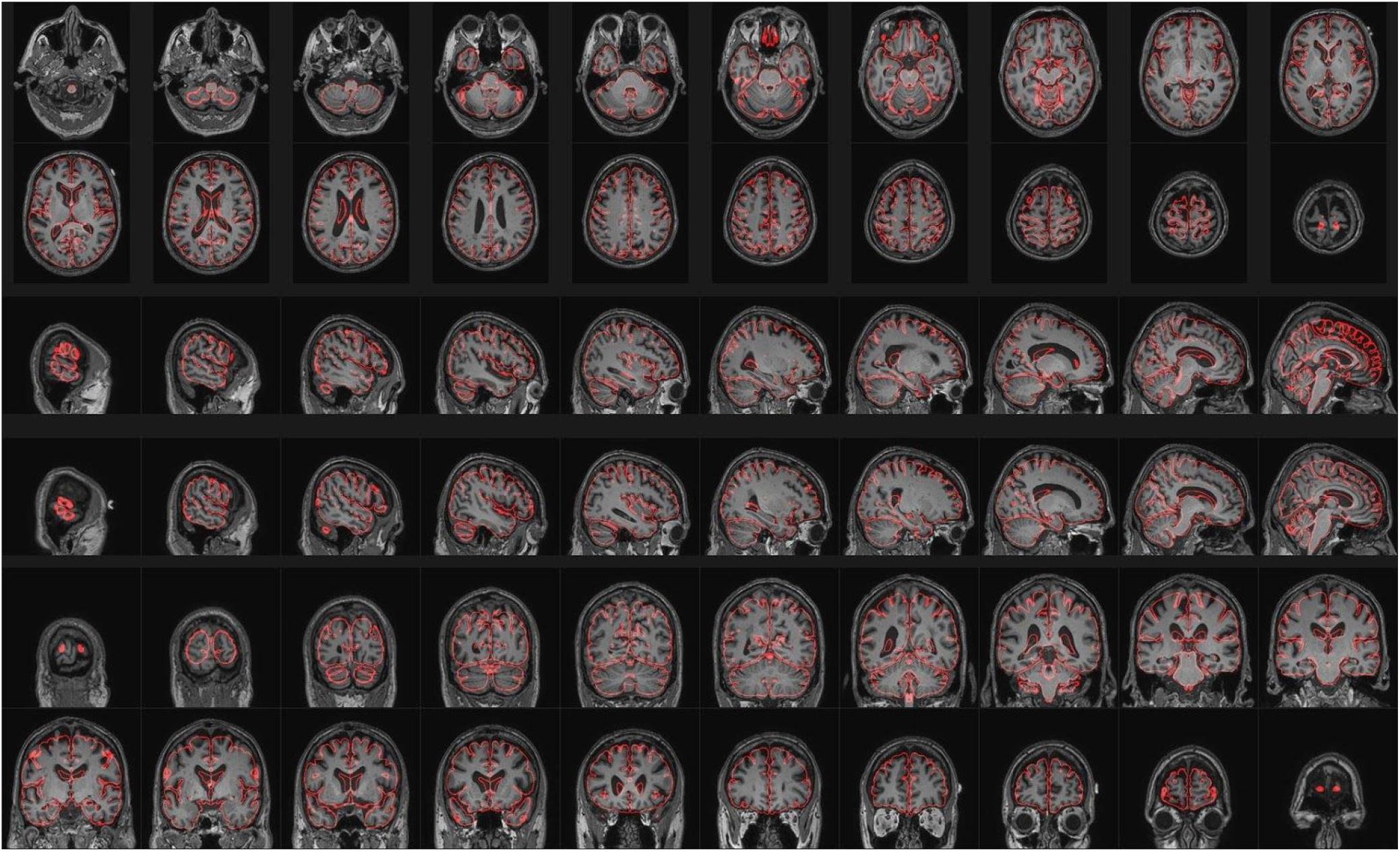
QC image for the human rater. Grayscale - one example subject’s MRI scan after registration in stereotaxic space, red line - outline of the MNI-ICBM152-2009c brain template.

Out of 57848 examples, 46231 (**79.9**%) were accepted (passed) and 11617 (**20.0**%) rejected (failed). To test reproducibility of the human rater results, a random subset consisting of 1000 examples were re-evaluated by the same rater, resulting in intra-rater dice kappa similarity of **0.96**, accuracy of **93**% and false-positive rate of **17.3**%.

### 2.4 Automatic registration quality control with deep neural network

We modified an existing deep neural network design for this task, reusing weights trained on the ImageNet database [9,10,11]. Instead of feeding a single image with different views showing various slices of the registered image as in the human quality control process, we used a stack of images that were created in the following fashion: (i) 3D MRI scan in the native space, without any preprocessing, was resampled to the MNI template space using the linear transformation matrix provided by registration algorithms; (ii) the whole range of the image intensities of the input file was mapped to 0-1 range; (iii) one Axial, one Sagittal and one Coronal slice was extracted from the middle of the registered 3D volume; (iv) the 2D images of slices were resampled to have 256 pixels in the longest dimension and then cropped around the central area to the 224×224 pixels.

Thus for each dataset, this image stack is used as input features for the DNN. Figure 2a shows an example of images corresponding to a scan that passed QC and figure 2b shows one that failed QC. Figure 2c shows a reference image corresponding to MNI template.

**Figure 2.**
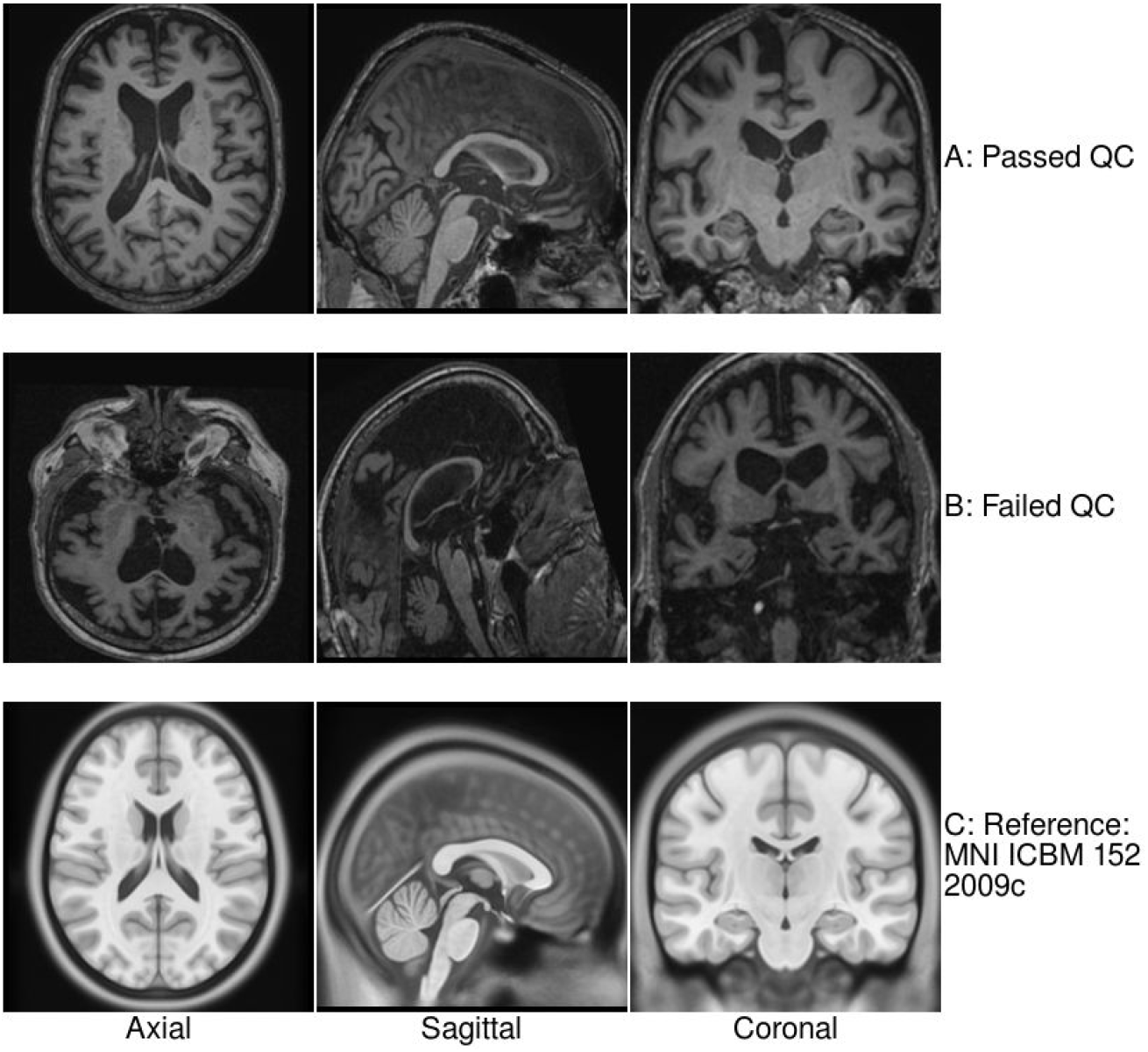
Images generated for automated QC script.

In order to transfer domain knowledge, we modified a DNN trained on the ImageNet dataset (DNN0) in the following fashion: (i) the input layer was altered to deal with grayscale images by collapsing the weight tensor along the dimension corresponding to input RGB dimension; (ii) the last few layers corresponding to high level features used for ImageNet classification were removed; (iii) each input feature from the image stack was processed sequentially using the same DNN0 layer; (iv) outputs of DNN0 layer corresponding to the three different images from the stack were concatenated and used as inputs to the last several layers, replicating the behavior of the layers originally removed from DNN0; (v) the final layer was modified to produce only two labels - pass or fail.

We also created an alternative scheme, where reference images extracted from MNI template were used as an additional set of features in the image stack. In this case, DNN0 was modified to accept 6 grayscale feature maps - combining those that were subject-specific, and the others that were fixed for all subjects (see Figure 2c). A simplified schematic representation design of the new DNN is shown in Figure 3.

**Figure 3.**
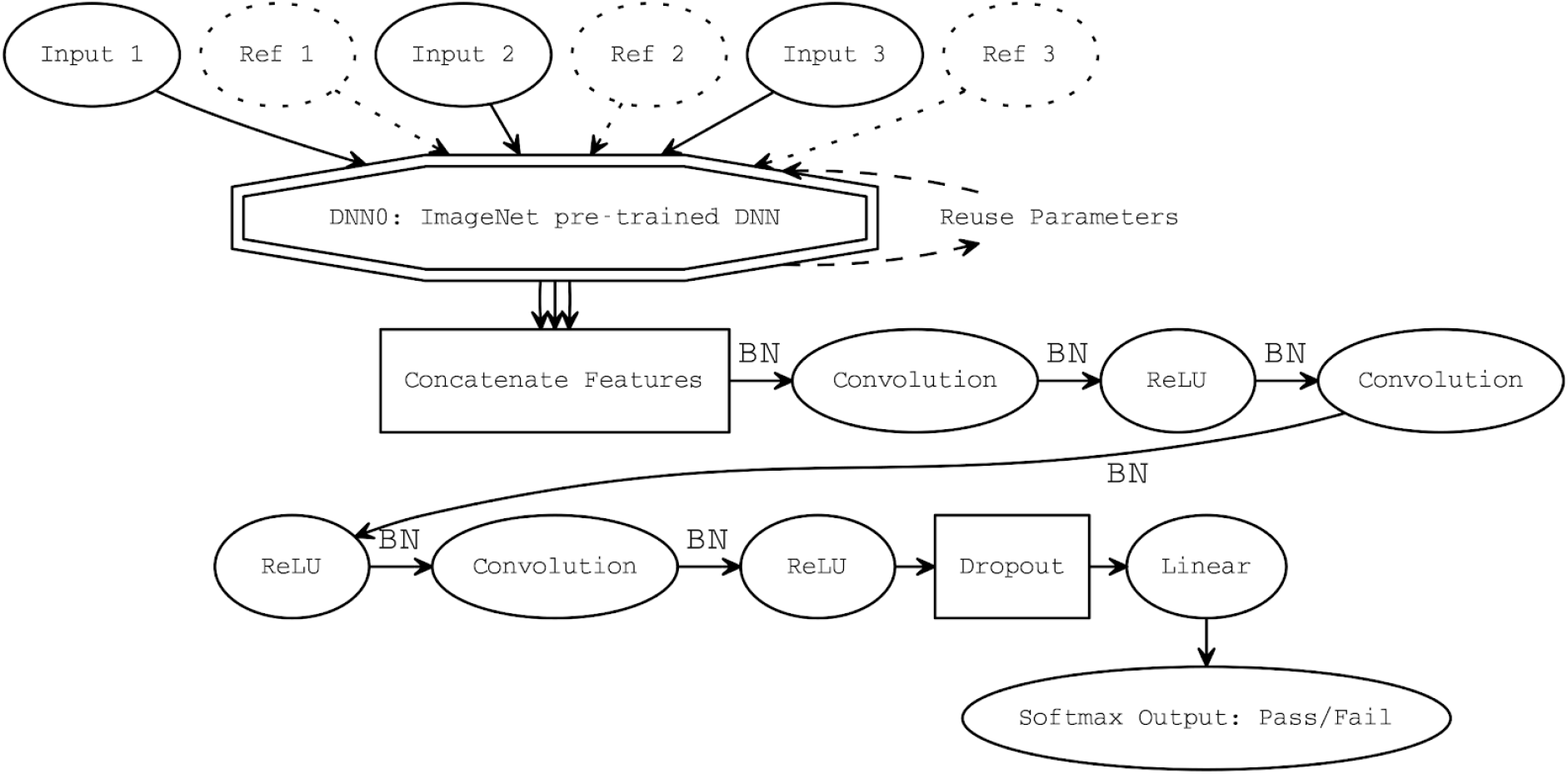
Overall DNN design, dotted nodes represent optional components; BN - batch normalization; ReLU – rectified linear unit

A cross-entropy loss function was used as objective function to train the DNN. The ADAM optimizer was used to train the network.

### 2.5 Network training

We used an 8-fold cross-validation scheme, where all original datasets (9693) are split into 8 equal partitions, at each round of cross-validation one corresponding registration results of each partition is used as “testing” dataset and the rest of data is used as “training” dataset. A small subset of “training” dataset (200 samples) was excluded from training and used for on-line validation for early stopping (“validation” dataset). The remaining training datasets were split into “pass” and “fail” subsets. At each iteration of the training, 32 samples from each subset were used to create a minibatch with balanced mix of pass/fail samples that was passed through DNN for training using the ADAM optimizer, two iterations of training were performed on each minibatch. Every 500 mini-batches, all “training” samples were re-shuffled.

An initial experiment was run with 20,000 mini-batches using one fold out of 8 to determine the optimal number of iterations based on the change of the false positive rate (FPR) of the “validation” dataset. See figure 4 for the progress of optimizer.

**Figure. 4.**
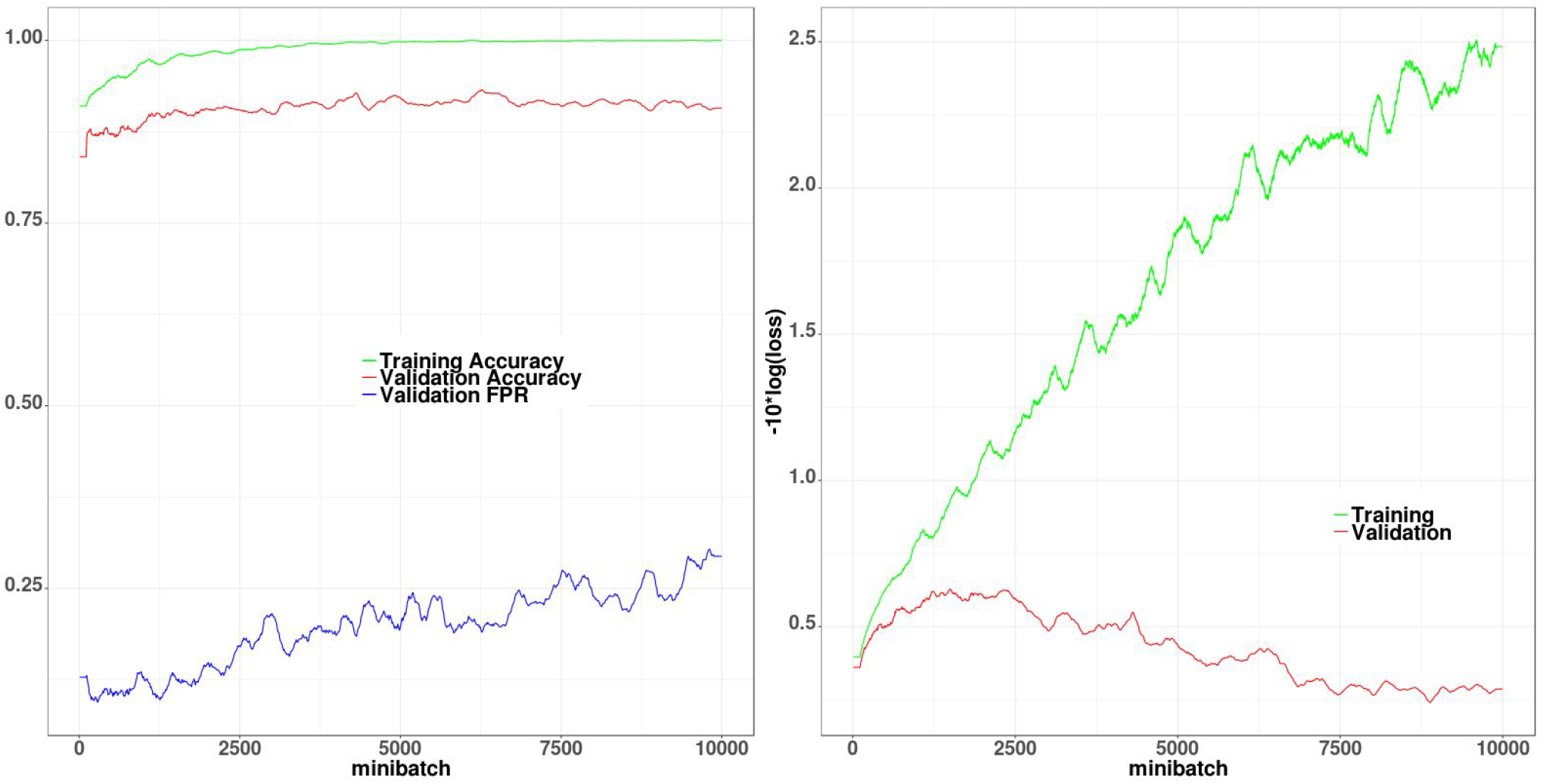
Progressions of DNN training for ResNet-18 without reference, values are smoothed with moving average with bandwidth of 200. Over-fitting is visible after approximately 1500 minibatches.

### 2.6 Silver standard and distance estimation

To characterize misregistration quantitatively, we created a “silver standard” transformation for each dataset, by averaging all transformations that passed manual QC, and then calculating the distance between each transformation and the “silver standard”, as defined in eq (1), where ROI_icbm_ is the bounding box of the brain ROI of the ICBM 152 2009c template [8].

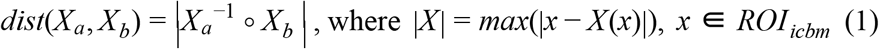

## 3 Results

Following the initial experiment with 20,000 mini-batches using one out of 8 folds, we observed over-fitting after approximately 1500 mini-batches, so the rest of experiments were conducted with 1500 minibatches.

Two types of pre-trained DNN were evaluated: Network-in-Network [6] and ResNet-18 [7, 11]. Two training schemes were tested - using only QC images (No Reference) and combining QC images with the images of the reference dataset: MNI template (With Reference). Resulting accuracy (acc), false-positive-rate (fpr), true-positive-rate (tpr) and area under receiver operating characteristic curve (auc) [21] are shown on Figure 5, the receiver operating characteristic curves with respect to the output of softmax layer are shown on Figure 6. The Resnet-18 DNN0 with reference images showed the best results in terms of false positive rate (**11%**) at the expense of the slight decrease of the overall accuracy (**88%**). Overall, all results are consistent with the performance of human rater (test-retest accuracy of **93%** and false-positive rate of **17%**).

**Figure 5.**
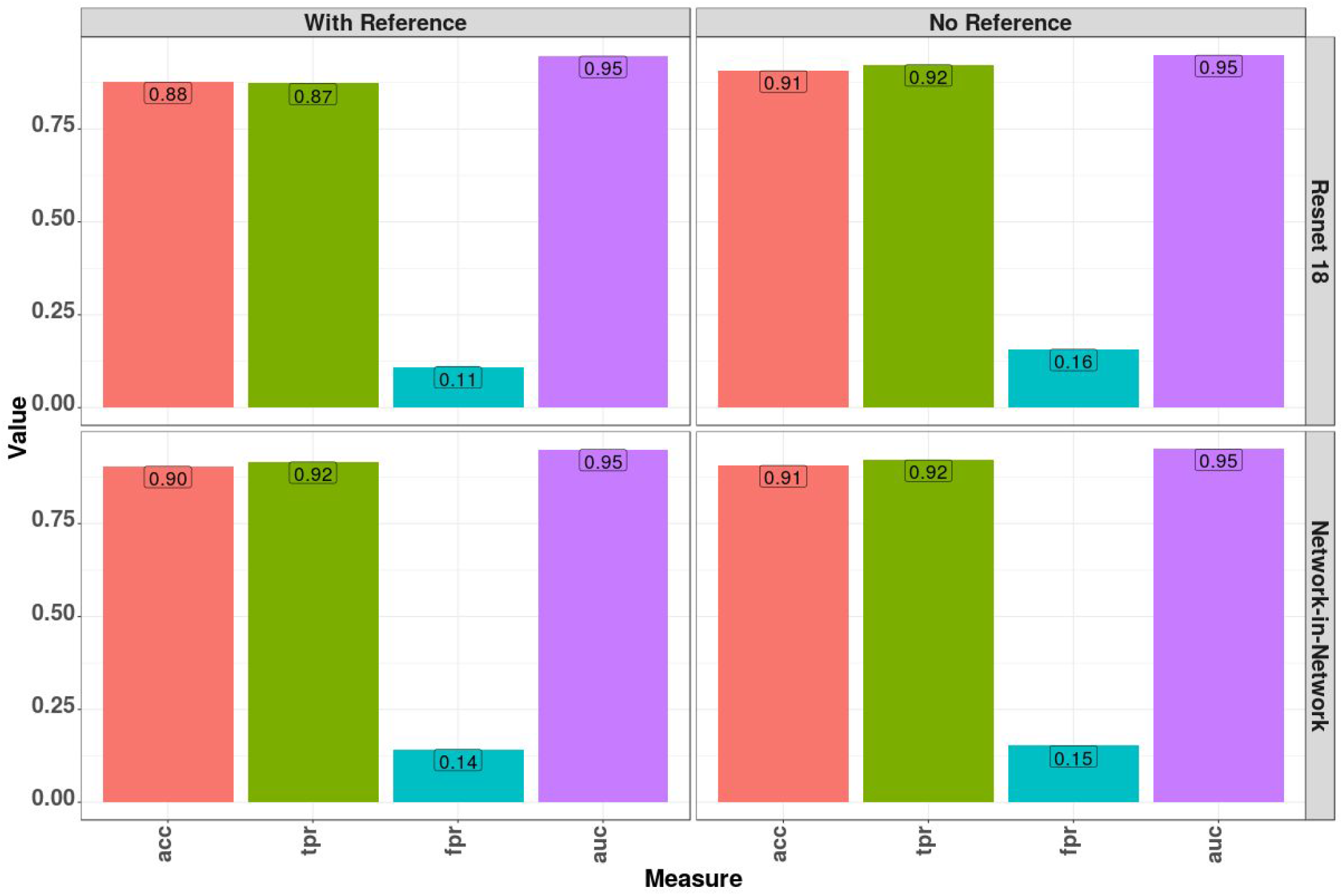
Performance of four tested combinations of methods: acc - accuracy; auc - area under the curve from ROC analysis; tpr - true positive rate; fpr - false positive rate.

**Figure 6.**
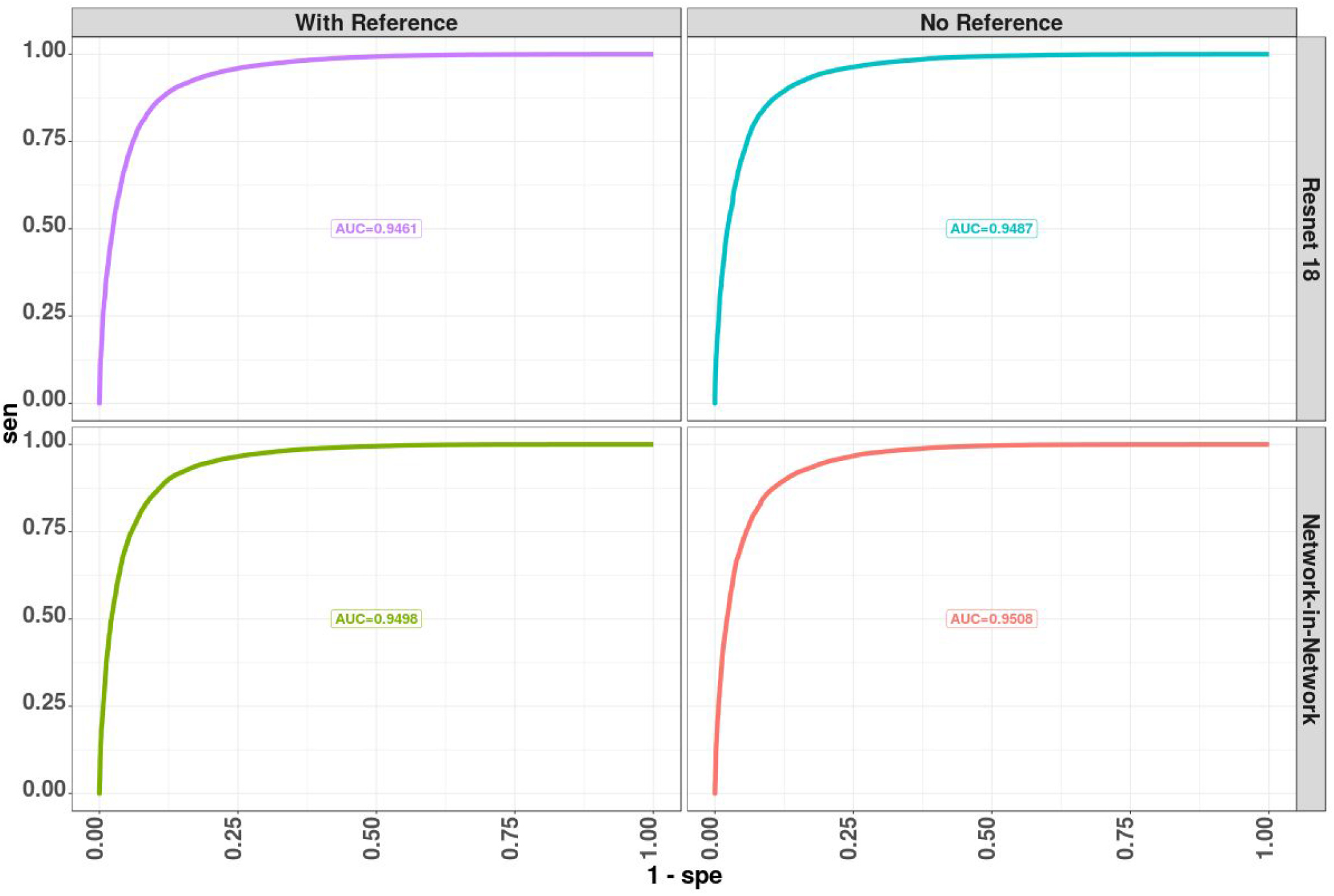
Receiver operating characteristic curves of each method, using softmax output of the final stage of the neural net.

If instead of using a threshold of 0.5 for the soft-max layer, a threshold of 0.95 is used to determine the samples that pass QC, the overall accuracy of all method would decrease (see Figure 7), and in this case, resnet-18 with reference would achieve a false-positive rate of 0.045 and the true-positive-rate decreases to 0.66.

**Figure 7.**
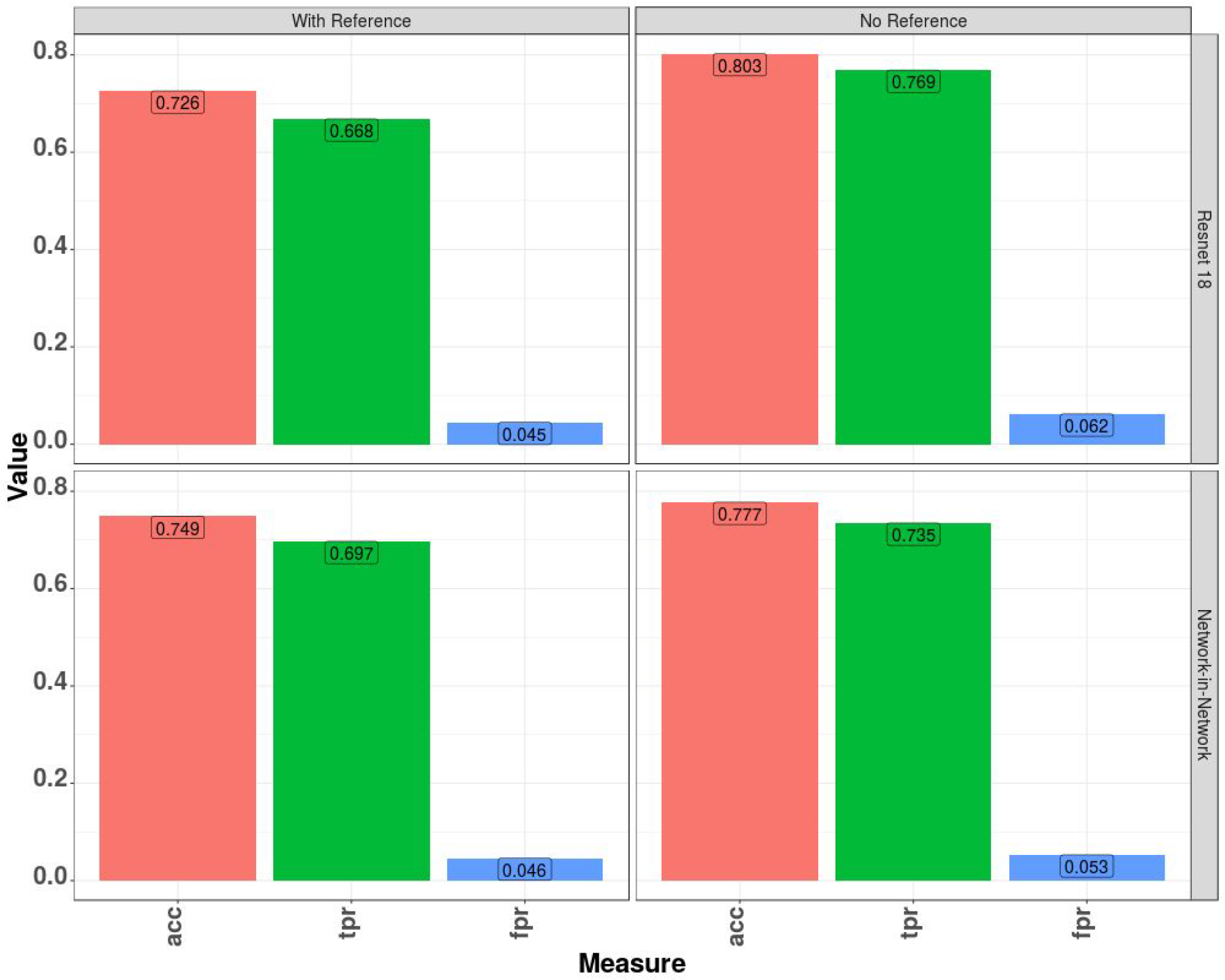
Performance of the methods, after thresholding softmax output at 0.95, instead of 0.5 to determine the pass/fail.

The distribution of the distances between a “silver standard” transformations and transformations estimated by each method is shown on Figure 8, the behaviour of the distances depending on the automated QC outcome is also shown on Figure 8.

**Figure 8.**
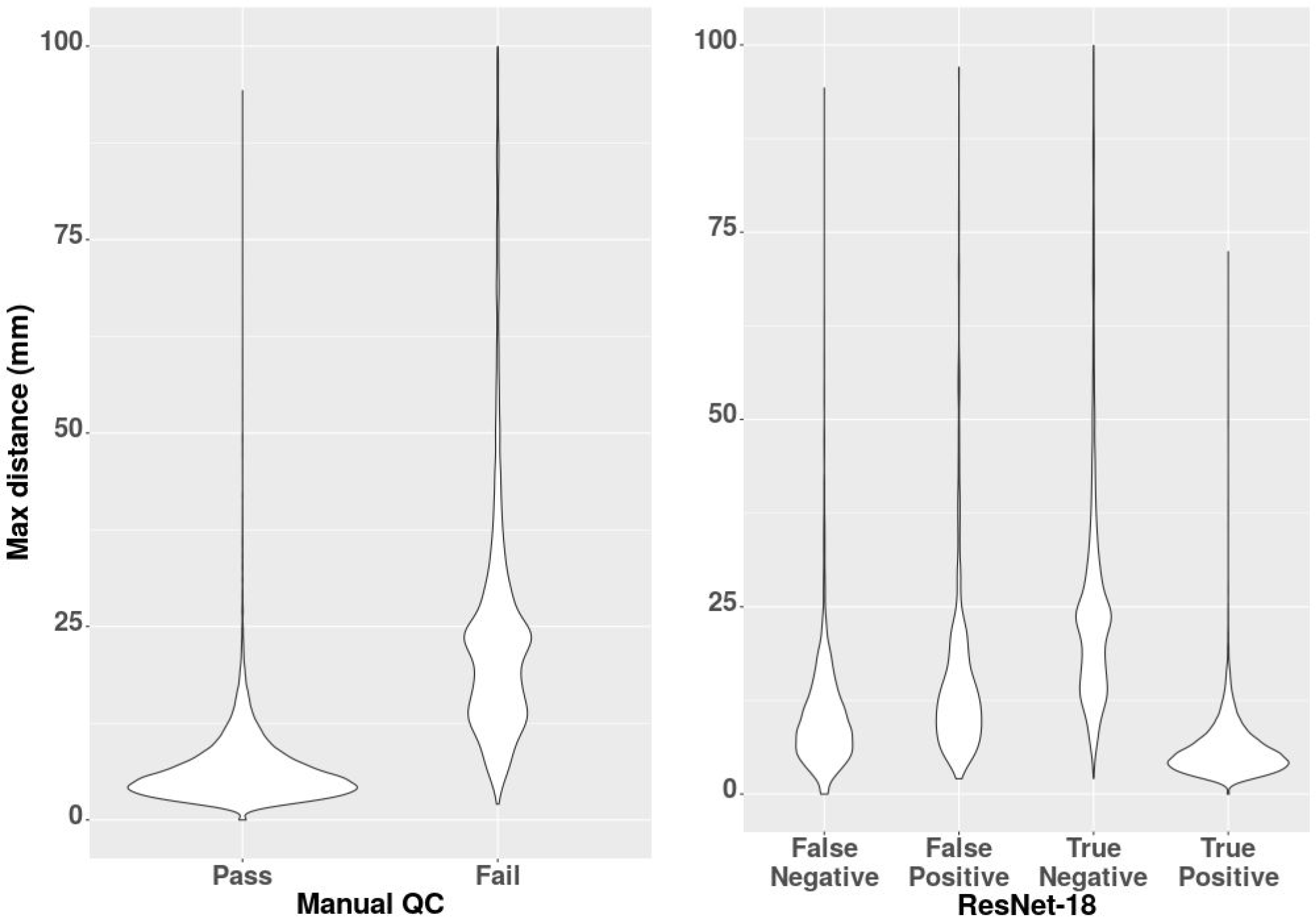
Distance from the “silver standard” in mm, for the manual QC results, and for the outcome of Resnet-18 method cross-validation.

## 4 Conclusions

We have demonstrated that it is possible to automate the task of manual quality control using a deep learning network with results comparable to the human rater.

This technique will save significant amount of human effort in processing large imaging databases and will increase reproducibility of results. And could be used for automatic testing of the robustness of image registration methods.

In addition, removing human effort from the registration QC process enables the users to rerun the registration methods with different settings and parameters to ensure a high acceptance rate for their cohort of interest. In other words, the proposed QC procedure can be incorporated into the process of the registration to enforce the method to repeat registration with different parameter settings until an accepted registration is obtained. A false negative (failing an acceptable registration) in this case would only increase registration time by forcing the method to repeat the process. Therefore, a lower tpr at the expense of a low fpr would be tolerable and lead to the overall improvement of the registration performance.

Unfortunately, the performance of the automated QC method is limited by the performance of the manual rater, that was available to perform training. To improve this method, it is possible to improve manual quality control and create better a “silver standard” as a reference. It would allow to simulate “failures” of registration and train network to clearly detect when mis-registration exceeds some pre-set threshold, but this approach is not necessary going to produce kind of errors that would naturally happen in considered registration techniques.

### 4.1 Implementation

Original implementation was done using torch library [22], we have re-implemented software in pytorch [23] since.

The source code of the method, implemented in torch and pytorch (python) and pre-trained neural network is available at https://github.com/vfonov/deep-qc

## 5 Acknowledgements

ADNI data is from the Alzheimer’s Disease Neuroimaging Initiative (ADNI) (National Institutes of Health Grant U01 AG024904). PPMI data was obtained from the Parkinsons Progression Markers Initiative (PPMI) database (www.ppmi-info.org/data). HCP data was obtained from the Human Connectome Project, WU-Minn Consortium (1U54MH091657). PREVENT-AD data were obtained from the Pre-symptomatic Evaluation of Novel or Experimental Treatments for Alzheimer’s Disease (PREVENT-AD, http://www.prevent-alzheimer.ca) program data release 3.0 (2016-11-30). Data collection and sharing for this project were supported by its sponsors, McGill University, the Fonds de Research du Québec - Santé, the Douglas Hospital Research Centre and Foundation, the Government of Canada, the Canadian Foundation for Innovation, the Levesque Foundation, and an unrestricted gift from Pfizer Canada. Private sector contributions are facilitated by the Development Office of the McGill University Faculty of Medicine and by the Douglas Hospital Research Centre Foundation (http://www.douglas.qc.ca/).

